# Mechanisms underlying the cooperation between loss of epithelial polarity and Notch signaling during neoplastic growth in *Drosophila*

**DOI:** 10.1101/2020.02.07.938621

**Authors:** Rémi Logeay, Charles Géminard, Patrice Lassus, Miriam Rodríguez-Vázquez, Diala Kantar, Lisa Héron-Milhavet, Bettina Fischer, Sarah J. Bray, Jacques Colinge, Alexandre Djiane

## Abstract

Aggressive neoplastic growth can be initiated by a limited number of genetic alterations, such as the well-established cooperation between loss of cell architecture and hyperactive signaling pathways. However, our understanding of how these different alterations interact and influence each other remains very incomplete. Using *Drosophila* paradigms of imaginal wing disc epithelial growth, we have monitored the changes in Notch pathway activity according to the polarity status of cells (*scrib* mutant). We show that the *scrib* mutation impacts the direct transcriptional output of the Notch pathway, without altering the global distribution of Su(H), the Notch dedicated transcription factor. The Notch-dependent neoplasms require however, the action of a group of transcription factors, similar to those previously identified for Ras/*scrib* neoplasm (namely AP-1, Stat92E, Ftz-F1, and bZIP factors), further suggesting the importance of this transcription factor network during neoplastic growth. Finally our work highlights some Notch/*scrib* specificities, in particular the role of the PAR domain containing bZIP transcription factor and Notch direct target Pdp1 for neoplastic growth.

## INTRODUCTION

Epithelial cells represent the basic unit of many organs. They are polarized along an apico-basal (A/B) axis, a feature critical for many aspects of their biology. A/B polarity is controlled by the asymmetric segregation of highly conserved protein complexes such as the Scrib/Dlg/Lgl complex (Bilder et al., 2003; Coopman and Djiane, 2016; St Johnston and Ahringer, 2010). The far-reaching effects of A/B polarity is epitomized by the observation that many tumors of epithelial origin exhibit impaired polarity, and that several viral oncoproteins target polarity complexes (Banks et al., 2012; Huang and Muthuswamy, 2010).

Studies in human cell lines and in animal models have also suggested a contributing role of polarity alterations to tumor formation. For instance, mutations in the baso-lateral determinant *SCRIB1* have been shown to control proliferation and invasion in MCF-10A human mammary cells (Cordenonsi et al., 2011). Similarly in *Drosophila, scrib*, *dlg* or *lgl* mutations, result in multilayered overgrowth of larval epithelial imaginal discs (Bilder et al., 2003; Bunker et al., 2015). However, this uncontrolled growth is at least partly achieved because larvae exhibiting *scrib* mutations fail to undergo proper metamorphosis and imaginal discs grow for an extended period. Indeed, *scrib* mutant cells actually grow slower than wild-type cells and are eliminated by wild-type neighbors (Cordero et al., 2010; Igaki et al., 2009, 2006; Ohsawa et al., 2011). Interestingly, this is reversed when additional mutations are introduced, such as overexpression of the BTB/POZ chromatin remodelers Abrupt or Chinmo (Doggett et al., 2015; Turkel et al., 2013) or the constitutive activation of signaling pathways (e.g. Ras or Notch), converting *scrib* mutant cells into aggressive, invasive and hyperproliferative cells (Brumby and Richardson, 2003; Pagliarini and Xu, 2003). Similar observations have been reported in mouse, where Notch or Ras activation and *Par3* depletion cooperate to generate aggressive neoplasms in mouse mammary glands (McCaffrey et al., 2012; Xue et al., 2013).

The Notch pathway is a highly conserved cell-signaling pathway mis-regulated in several cancers (Ntziachristos et al., 2014; Ranganathan et al., 2011). Upon activation, Notch receptors undergo two proteolytic cleavages to release their intra-cellular domain or NICD, which enters the nucleus, binds to the Notch pathway specific transcription factor CSL (Rbpj in mammals; Suppressor of Hairless, Su(H) in *Drosophila*), and converts it from a repressor to an activator to turn on the transcription of specific target genes (Bray, 2016). These Notch direct target genes differ depending on cell type and account for the variety of outcomes triggered by Notch activity. Increased Notch activity has been associated with several epithelial cancers such as non-small-cell lung carcinomas (Maraver et al., 2012; Ntziachristos et al., 2014), but in animal models, the sole increase in Notch activity either promotes differentiation or only results in benign over-proliferation (hyperplasia) (Brumby and Richardson, 2003; Djiane et al., 2013; Fre et al., 2005; Ho et al., 2015; McCaffrey et al., 2012). However, as mentioned previously, Notch pathway activation cooperates with loss of polarity to generate invasive neoplasms (Brumby and Richardson, 2003; Ho et al., 2015; McCaffrey et al., 2012; Pagliarini and Xu, 2003).

So, while the cooperation between loss of cell architecture and hyperactive signaling pathways is well established, the underlying mechanisms remain poorly understood. It could merely reflect an additive effect where the consequences of both events combine. Alternatively, it could indicate a more profound integration within epithelial cells where these two events impact on each other to generate unique new behaviors. Using *Drosophila* paradigms of imaginal wing disc epithelial growth, we have monitored the changes in Notch pathway activity according to the polarity status of cells and show that epithelial polarity changes directly impact the transcriptional output of the Notch pathway. We further provide evidence that this Notch redirection is not mediated by new genomic binding regions for Su(H), but relies on the cooperation with Su(H) of a combination of transcription factors, such as Stat and basic leucine zipper (bZIP), whose activity is triggered in response to JNK signaling during polarity loss, extending earlier reports on the cooperation between oncogenic Ras and polarity loss (Atkins et al., 2016; Davie et al., 2015; Külshammer et al., 2015; Uhlirova and Bohmann, 2006). Our work highlights in particular the role of the PAR domain containing bZIP transcription factor Pdp1 for Notch-driven neoplastic growth.

## RESULTS

### Notch activation and scrib mutation cooperate to promote neoplastic growth

In order to gain insights into the mechanisms underlying neoplastic growth, we first characterized the effects of Notch activation and *scrib* mutation mediated epithelial polarity impairment on wing disc growth. Using precisely controlled *Drosophila* larvae culture conditions (crowding and timing), we compared the phenotypes of wild-type (WT), Nicd overexpressing (N), *scrib* mutant (S), and Nicd overexpressing and *scrib* mutant (NS) 3rd instar wing imaginal discs at 6 days after egg laying at 25C. These different paradigms are shown in Figure 1A-D. For clarity, in all figures N will be shown in green, S in red, and NS in blue. Reproducing our previous observations (Djiane et al., 2013), N discs overgrew compared to WT, but remained as monolayered epithelia, and represent a paradigm of hyperplastic-like growth (Fig. 1A&B). S discs were smaller than WT, but grew as unstratified mass of cells. These discs however showed an extensive expression of the JNK signaling target Mmp1 (Fig. 1C), a metallo-protease implicated in the digestion of the extracellular matrix, indicative that *scrib-*cells activate JNK signaling and are prone to invasiveness (Igaki et al., 2006; Uhlirova and Bohmann, 2006). It is noteworthy that S larvae did not pupariate and if left to grow for longer, the S discs ultimately developed as massive overgrowths with very disrupted epithelial polarity, that invaded and fused with neighboring tissues such as other discs (Bilder et al., 2003). Strikingly, NS discs combined aspects of N and S discs. They were overgrown like N discs, but also expressed high levels of Mmp1 like S discs (Fig. 1D). These discs grew as multilayered tissues and were able to invade the surrounding tissues such as haltere discs, and represent therefore a paradigm for neoplastic-like growth.

**Figure 1.**
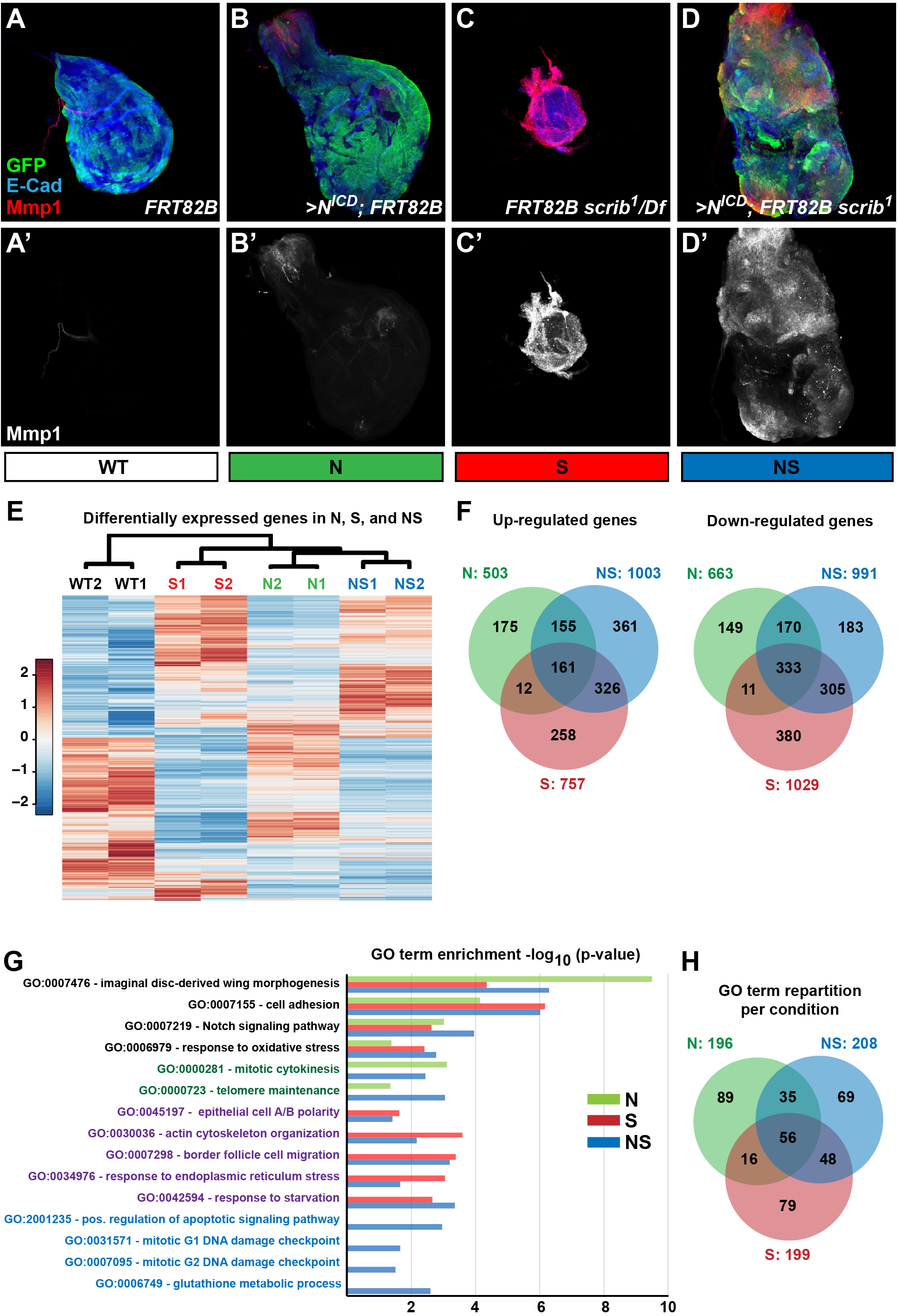
Notch-based growth paradigms in *Drosophila* wing discs. **A-D.** 3rd instar wing imaginal discs at precisely 5 days after egg-laying either wild-type (WT; A), overexpressing activated Notch (N; B), mutant for *scrib* (S; C), or combining overexpressed Notch and *scrib* mutation (NS; D) and marked for E-Cad (blue) and Mmp1 (red). **A,B&D.** MARCM clones (positively marked by GFP; green) of the indicated genotypes: expressing only GFP (A; WT), expressing Nicd & GFP (B; N), and expressing Nicd & GFP and mutant for *scrib* (C; NS). **C.** Discs fully mutant for *scrib*. **E-F.** Differentially expressed genes as compared to WT in the different growth paradigms N (green), S (red), and NS (blue) identified by RNA-Seq. This color code, green for N, red for S, and blue for NS is used in all figures. **E.** Heatmap of gene expressions after unsupervised clustering. **F.** Venn diagram of up-regulated and down-regulated genes in N, S, and NS. **G.** Enrichment diagram as measured by adjusted p-value for selected GO terms and represented as bars for N (green), S (red), and NS (blue). GO terms color reflect whether they are shared or specific: shared by all (black), common N&NS (dark green: mix of green and blue), common S&NS (purple: mix of red and blue). **H.** Venn diagram showing the domains of overlap of GO terms identified (significantly enriched) in N, S, and NS.

### Neoplastic and hyperplastic discs have different transcriptomes

The context of the *scribble* mutation converts the Notch-based *Drosophila* wing disc hyperplasia into a paradigm of neoplastic growth. While the cooperation between activated Ras and *scrib* mutations has been studied extensively, mainly in the eye imaginal disc (Atkins et al., 2016; Cordero et al., 2010; Davie et al., 2015; Igaki et al., 2009; Katheder et al., 2017; Pagliarini and Xu, 2003; Toggweiler et al., 2016; Wu et al., 2010), less attention has been given to the cooperation between polarity loss and other activated pathways, such as Notch or Hedgehog (Brumby and Richardson, 2003; Pagliarini and Xu, 2003), preventing an evaluation of how general the studies on Ras signaling are to neoplasia development.

First, we compared the RNA-Seq transcriptomes of the different genetic conditions, to identify the differently expressed genes in N, S, and NS compared to WT controls (using DESeq with adjusted p-value for multiple testing <0.05; Fig. 1E&F and Supplemental Table S1; (Anders and Huber, 2010). The numbers were broadly similar in the different conditions, N (503 up; 663 dw), S (757 up; 1029 dw), and NS (1003 up; 991 dw). Semi-quantitative qRT-PCR validated in a subset of genes the transcriptional changes. For instance, *E2f1*, *Sdr*, and *mxc*, were activated only in N discs, while *Act87E* and *Wnt10*, were activated only in NS. In addition, *Ets21C*, *ftz-f1*, and *Atf3* were up-regulated in all conditions (Supplemental Fig. S1A). Comparing these data with previously published transcriptomes on similar or related genetic backgrounds revealed significant overlap, validating our experimental approaches. For instance, 174/503 up-regulated, and 285/663 down-regulated genes in N were also detected in our previous analysis using dual color differential expression arrays (significant overlap p=1.11e-273, hypergeometric test; (Djiane et al., 2013). Similarly, 676/757 up-regulated genes in S were identified in a previous analysis of *scrib* depleted discs (significant overlap p=4.90e-193; (Bunker et al., 2015).

GO term analysis (p-value < 0.05) confirmed that, as expected from their genetic composition, the N and NS transcriptomes had over-representation for genes in the Notch signaling pathway (GO:0007219) while the NS and S transcriptomes for genes affected by changes in A/B polarity (GO:0045197; GO0019991) (Fig. 1G). This analysis revealed potential common behaviors shared by N and NS, in particular signs of increased proliferation, consistent with the overgrowth phenotypes (e.g. “mitotic cytokinesis” and “mitotic spindle organization”; GO:0000281; GO:0007052), or behaviors shared between NS and S, such as those related to cell migration (e.g. “border follicle cell migration”; GO:0007298) or cellular stress (e.g. “response to starvation” GO:0042594; “response to endoplasmic reticulum stress” GO:0034976) (Fig. 1G). Notably several GO categories were enriched specifically in NS, including “mitotic G1/G2 DNA damage checkpoint” (GO:0031571/GO:0007095), or gluthatione metabolic process (GO:0006749). These results argue that the combined Notch activation and polarity loss promoted the emergence of new cell behaviors and responses, in particular related to DNA damage responses (Fig. 1G).

### Hyperplasia and neoplasia harbor different Notch direct target networks

This raises the question of how the defects in *Notch* and in *scrib* cooperate to produce these transcriptional consequences. One particularly critical aspect is how the Notch pathway is affected by the *scrib* mutation. Many signaling pathways, such as Ras, branch and act through several transcription factors and/or combine nuclear and cytoplasmic responses, making the analysis of how they are affected by the *scrib* mutation complicated. Since the Notch pathway is extremely direct and since the major, if not unique, output of Notch signaling is transcriptional activation (Bray, 2016), it offers a unique opportunity to investigate how the *scrib* mutation could potentially affect Notch signaling in the NS cooperation. We thus decided to monitor the genes directly activated by Notch and study whether the Notch direct programme (Notch Direct Targets, NDTs) is affected by the *scrib* mutation.

Genes that are directly regulated by Notch (NDTs) should have the transcription complex, containing Su(H) bound at their regulatory regions (Djiane et al., 2013). To identify potential NDTs in N and NS, we thus monitored the genomic regions occupied by the Su(H) transcription factor by genome-wide Chromatin Immuno-Precipitation (ChIP) (Fig. 2A; Supplemental Table S2). These overlapped significantly with our previous analysis of Notch induced overgrowth, suggesting that we have captured all the robust regions of Su(H) enrichment.

**Figure 2.**
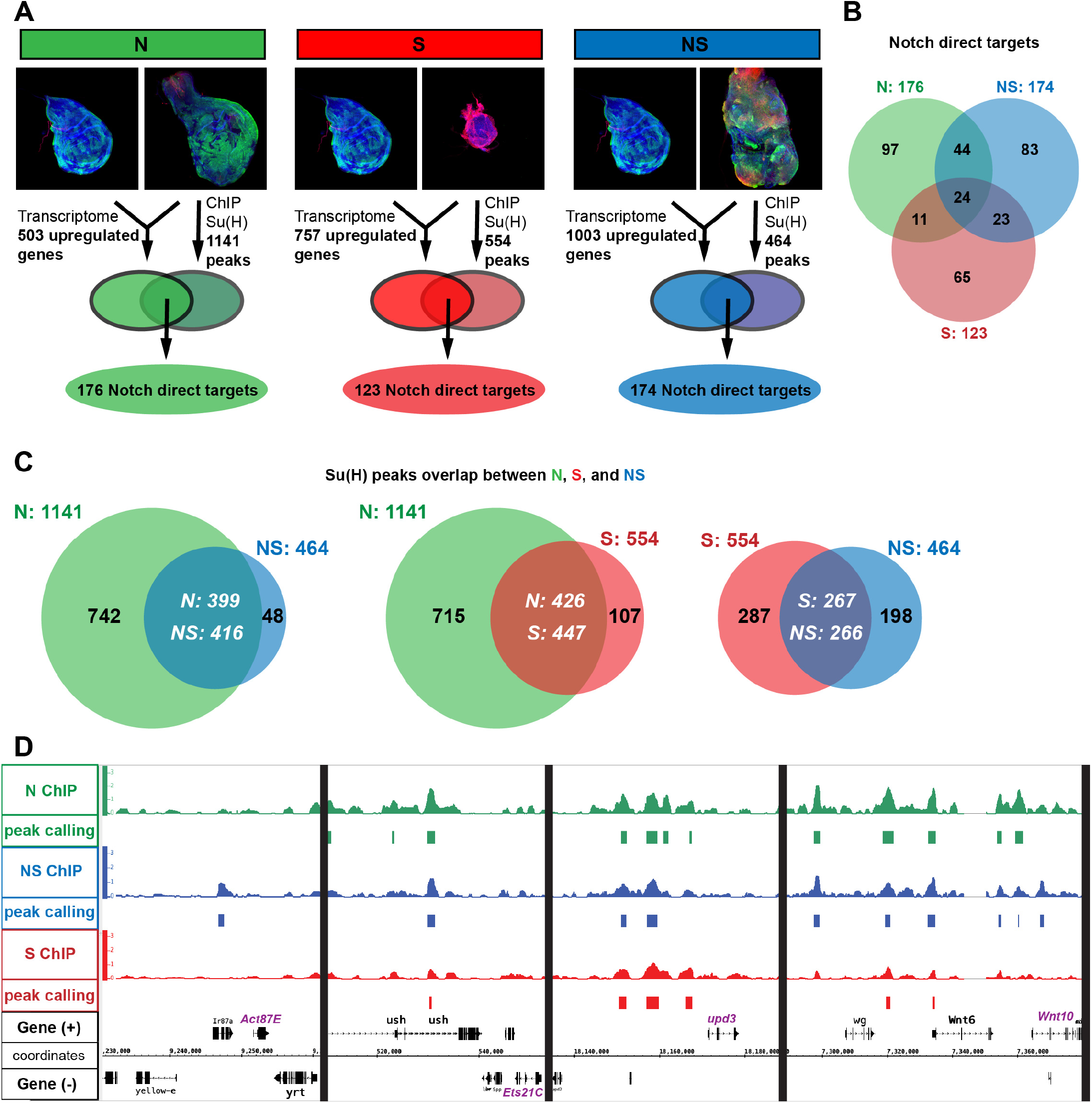
Polarity loss redirects the transcriptional output of Notch during neoplastic growth. **A.** Experimental set-up to identify the Notch Direct Targets genes (NDTs): Genes up-regulated in N, S, or NS (Transcriptomic), and located within 20kb of a Su(H) binding site (ChIP). **B.** Venn diagram showing the NDTs overlap in N, S, and NS, showing core Notch responses, but also significant condition specific NDTs. **C.** Overlap of the Su(H) binding sites identified by ChIP in N, S, and NS, showing that almost all S and NS Su(H) peaks are also found in N. The overlap is shown in white. Numbers are slightly different because in the Su(H) peaks calling protocol, in some rare cases, some peaks can be split between conditions where one peak in one condition would overlap with two peaks in the other. **D.** Genome Viewer snapshots of several NDTs (shown in purple) such as the NS specific *Act87E* and *Wnt10* (but also *yellow-e* and *Ir87a*), the common *Ets21C*, and the N/NS NDT *upd3*. For each condition, the Su(H) ChIP enrichment is shown in the upper lane, and the Su(H) peaks identified are represented by the blocks underneath.

Strikingly, there was a strong overlap in the Su(H) bound regions between N and NS conditions: almost all NS Su(H) peaks (416 out of a total of 464) overlap with peaks present in N discs. This implies that the vast majority of Su(H) binding in NS were also present in N (Fig. 2C & S1C). The overlap was also important between N and S peaks (447/554 S peaks overlapping with N peaks; Fig. 2C & S1C). These results suggests that in NS neoplastic discs, a minority of Su(H) peaks represent new binding regions compared to hyperplastic N discs, and that the new NS behaviors are not the consequence of general redistribution of the Notch specific transcription factor Su(H).

In order to estimate the programs specifically activated by Notch in N and NS, we then intersected the transcriptomic data with the Su(H) ChIP data, considering that upregulated genes located within 20kb of a Su(H) peak were likely NDTs. Using this approach, we identified similar numbers of NDTs in N (176) and NS (174) (Fig. 2A&B, Supplemental Table S3). Again, there was substantial overlap with previous data, with 64/176 NDTs in the N condition being identified in our previous study (significant overlap p=5.89e-96, hypergeometric test; (Djiane et al., 2013). When the N and NS scenarios were compared, 68 genes were common to both and thus represent core NDTs in the wing disc overgrowth. However, a significant proportion of NDTs appeared specific for each condition: 108 for N, 106 for NS. Amongst the 106 NS-specific NDTs, 23 were also NDTs in S. Indeed, affecting only polarity with the *scrib* mutation already alters the Notch programme (123 NDTs in S). But the difference between N and NS cannot just be explained by this S contribution as it concerns only 23 NDTs, and taking all comparisons into account, 83 genes appear as true NS-specific NDTs (Fig. 2B). Only a minority were associated with new Su(H) binding regions: around the *87E* locus (*yellow-e3, yellow-e, Ir87a*, and *Act87E*), and the *94A* locus (*CG18596, CG7059, CG13857*, and *CG13850*), but also next to the *vito*, *cdi*, and *REG* genes (see Supplemental Table S3).

These results indicate that while a core Notch response can be identified in overgrowing wing discs (68 genes; Fig. 2B), the loss of polarity affected the direct transcriptional output of the Notch signaling pathway: 108 NDTs specific to N were lost, while 83 NDTs specific to NS were gained. Importantly these changes in the Notch program are only marginally mediated by a redeployment of the Su(H) transcription factor: 11/83 NS NDTs correspond to NS-specific Su(H) enrichment.

### Neoplastic overgrowth is not mediated by the new NS-specific Su(H) regions

Using the NDT datasets, we sought to identify the factors that are required for the transition from N hyperplastic to NS neoplastic growth.

First, we decided to investigate the contribution of the DNA damage response. Indeed several NS-specific NDTs are implicated in DNA damage response (e.g. *p53*, *His2Av*) and the GO analysis highlighted categories specific to NS related to “response to oxidative stress” (GO:0006979), “cellular response to gamma radiation” (GO:0071480), and “DNA damage checkpoints” (GO:0031571/0007095). We thus asked whether interfering with such pathways could block the growth and invasiveness of NS tissues. To perform these genetic tests, we generated a stable fly line which overexpressed *Nicd* and *scribRNAi* together with a *GFP* marker under the *Bx-Gal4* driver (driving expression in the pouch of the larval wing discs; *Bx>NS*), and monitored the size of the overgrowth (GFP positive tissue), and its invasiveness potential (Mmp1 expressing cells; Fig. 4A).

Blocking the oxidative stress response by overexpressing the Reactive Oxygen Species (ROS) sponge CAT and SOD, did not have any significant effect on the NS overgrowth or the expression of Mmp1 (Fig. 4E&F). Similarly, expression of RNAi or dominant negative forms of the severe DNA damage major effector and NS specific NDT *p53* could not modify the NS overgrowth phenotype (Fig. 4E&F). While we cannot exclude that the tools used here were not strong enough, these results suggest that even though activated in NS tissues, oxidative stress and p53-mediated DNA damage responses were either not required to fuel NS growth, or that they could compensate for each other converging ultimately on an as yet unidentified core response promoting NS growth.

Second, we turned our attention to the NS specific NDTs associated with unique Su(H) binding. Indeed, even though the emergence of new Su(H) binding is not the main mechanism driving the NS-specific Notch programme, it remains possible that these loci and the genes associated are functionally important for the NS neoplastic behavior. We thus asked whether interfering with these 11 specific NS NDTs could block the growth and invasiveness. Knocking down by RNAi the expression of the different genes associated with the *87E* locus (*yellow-e3, yellow-e, Ir87a*, and *Act87E*), and with the *94A* locus (*CG18596, CG7059, CG13857*, and *CG13850*) did not have any significant effect on the NS overgrowth. Similarly, we could not detect any change in NS tumor overgrowth after RNAi-mediated knock-down for *cdi*, *REG*, and *vito* (data not shown). It should be noted however that for several genes only one RNAi could be tested with the potential caveat of insufficient knock-down efficiencies (Xia et al., 2021). However, our results suggest that these NS-specific Su(H) NDTs are not required, at least individually, for NS overgrowth. But even though not strictly required, their overexpression, might still contribute, redundantly with other factors, to the overall NS neoplastic behaviors. Amongst these particular NS NDTs, *Act87E*, associated with the new Su(H) peak at locus *87E* (Fig. 2D) was the most robustly up-regulated in NS (see Supplemental Table S1 & Fig. S1A). When overexpressed in wild-type wing disc, or in combination with activated Notch, Act87E led to robust expression of the metalloprotease and JNK target Mmp1, and cell delamination. Act87E also induced robust expression of the effector caspase Dcp-1 caspase (Supplemental Fig. S2). *Act87E*, might thus represent a stress gene activated in NS initiating cell delamination and ultimately cell death, but whose role is not necessary and/or redundant with other NS-activated genes.

### Identification of the transcriptional networks in the different growth paradigms

Taken together our results indicate that even though the *scrib* mutation was able to change the transcriptional output of the Notch pathway, the difference between N and NS is not due to a redeployment of Su(H) to activate new NS-specific genes. Other factors brought upon the *scrib* mutation must thus influence gene expression.

We thus sought to identify transcriptional factors that could account for the cooperation between Notch and polarity loss. We used the iRegulon software (Janky et al., 2014; Verfaillie et al., 2015) to identify the transcription factors likely co-regulating the genes identified in N, S, or NS. Previous usage of iRegulon on *RasV12/scrib-*overgrown 3rd instar larval discs, highlighted an “oncogenic module” comprising of the Hippo pathway terminal effectors Yki/Sd, the JNK pathway regulated AP-1 factors (in particular Atf3, Kay, and CEBPG), the Jak/Stat pathway, Myc, Crp, and Ftz-F1, (Atkins et al., 2016; Davie et al., 2015; Külshammer et al., 2015).

Implementing iRegulon on our step-wise Notch-based paradigms allowed us (i) to identify transcriptional modules unique or shared between polarity loss only (S), proliferation only (N), and proliferation plus invasiveness (NS), and (ii) to assess the conservation of the “oncogenic module” identified previously with Ras in a Notch-driven neoplastic paradigm. We performed these analyses feeding iRegulon either with the lists of up-regulated genes in N, S, and NS (Fig. 3 & Supplemental Table S4) or with the lists of NDTs (Fig. S3 & Supplemental Table S5). We decided to focus our analyses on the up-regulated genes here since the overexpressed Nicd triggers transcriptional activation (Bray, 2016) and down-regulated genes might thus represent very indirect effects of the cooperation. Modules identified are presented as Venn diagrams (Fig. 3&S3) highlighting the common and specific transcription factor modules that were identified as likely master regulators of the transcriptional changes in the different conditions. Feeding either up-regulated genes (Fig. 3) or NDTs (Fig. S3) identified similar modules indicating that the Notch pathway condition-specific transcription programs are mediated, at least in part, by transcription factors that broadly affect the whole transcriptome. This is likely also true for other neoplastic paradigms such as Ras, even though this remains to be established. Importantly, iRegulon identified the Notch pathway dedicated transcription factor Su(H) in the N and NS transcriptomes.

**Figure 3.**
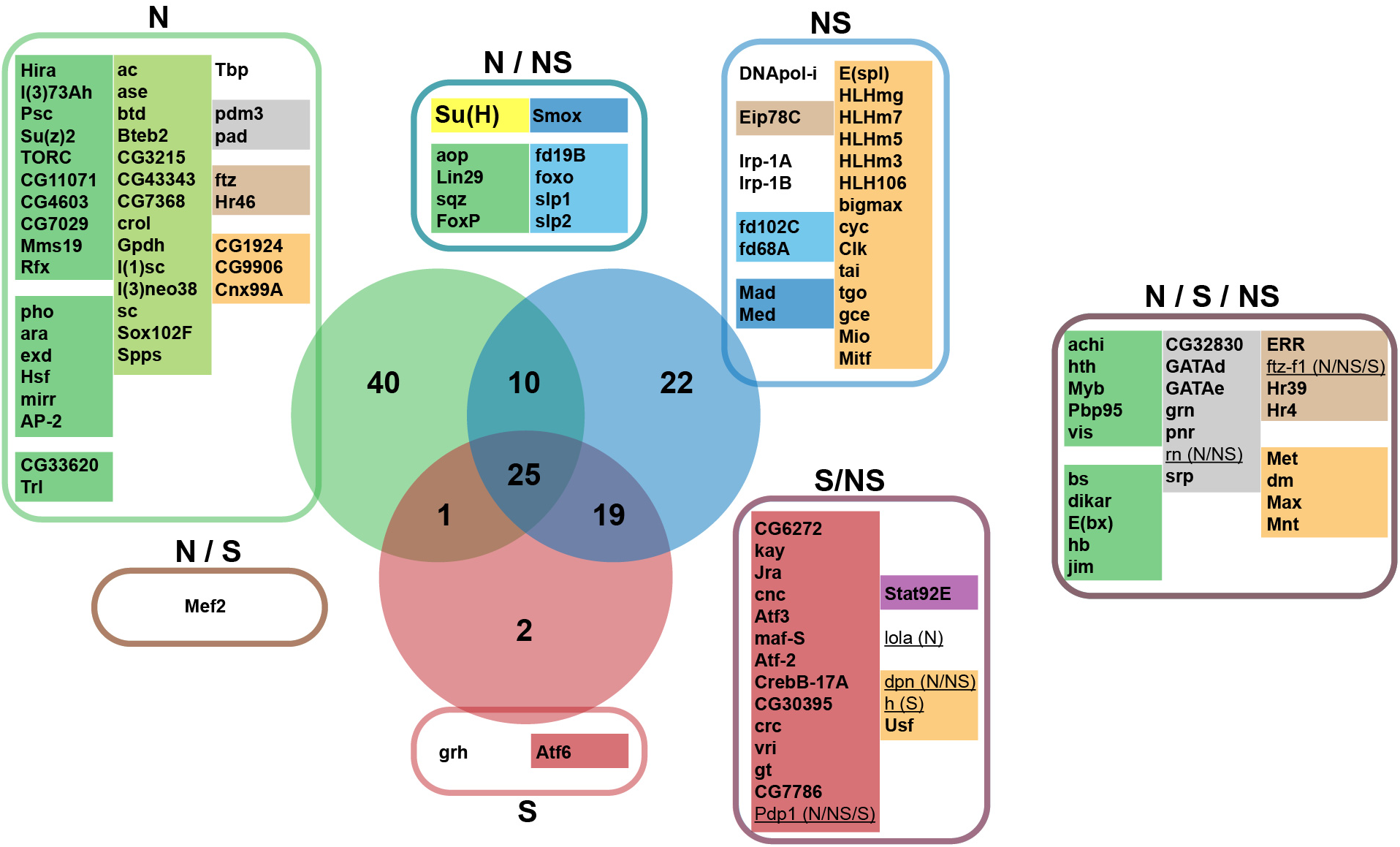
Identification of potential transcriptional modules mediating N, S, and NS growth. Venn diagram of significant transcription factors (TFs) identified by iRegulon as potential key mediators for the expression of the N (in the green circle), S (red circle), and NS (blue circle) up-regulated genes. Fed with lists of co-regulated genes, and analyzing the genomic features in the vicinity of the transcription start sites of these genes, iRegulon identifies potential groups of TFs and DNA-binding factors, that are enriched in the dataset of regulatory sequences, and could thus represent potential mediators of the N, S, and NS transcriptomes. For each condition, TFs were identified as part of “regulons” or group of TFs that could potentially together regulate the expression of subsets of the transcriptome. Taking only the TFs corresponding to the significant regulons identified in N, S, and NS, the shared and unique potentially regulating TFs are here presented in the different colored boxes. In each box, TFs were then grouped according to their molecular family (suggesting pretty similar binding motifs on the DNA), and color-coded: for instance, bZIP TFs in light maroon, or the nuclear receptors in brown. The molecular family color code was not respected for the green and light green groups which correspond to very diverse “regulons” that appear linked to epigenetic chromatin regulators/remodelers. For the complete list of regulons identified in each condition, and the nature of TFs and DNA binding proteins in each regulon, see Supplemental table S4). Numbers in the Venn diagram represent the number of TFs identified. TFs that are also NDTs and which could participate in feedforward loops are underlined (with their NDT condition in parentheses). See also Figure S3 for the iRegulon analyses of the NDTs, and the detailed lists of both iRegulon analyses in Supplemental Table S5.

First, focusing on NS, which most resembles the *RasV12/scrib-*paradigms, our analysis identified the same major nodes and the transcription factors associated with the “oncogenic module” as described previously for the *RasV12/scrib-*models:

- AP-1 basic Leucine Zipper factors related to stress kinase signaling
- Stat92E of the Jak/Stat pathway
- Ftz-F1 nuclear receptor
- basic Helix-Loop-Helix factors of the Myc family.

In NS transcriptome, we also identified a contribution of the E(spl) bHLH transcriptional repressors. *E(spl)-HLH* genes are canonical Notch targets, and they are robustly up-regulated in N and NS, in particular *E(spl)mγ-HLH*.

The iRegulon analyses also suggested that the AP-1 bZIP, and the Stat92E signatures in NS are contributed by S, since they are also detected in the S transcriptomes, while the Su(H) signature is contributed by N. Finally, iRegulon identified a signature for the Polycomb chromatin silencers specifically in N. Such factors include Pho, a zinc finger protein which binds to Polycomb Responsive Elements (PREs) and recruits Polycomb complexes, and the three Polycomb Repressor Complex 1 components Psc, Su(z)2, and l(3)73Ah. Recently, PRC1 has been associated with specific and unexpected transcriptional activation at larval stages, raising the possibility that in N, such genes, normally repressed at embryonic stages, become active (Loubiere et al., 2020). However, the exact contribution of Polycomb factors, and whether they are actually involved in gene activation upon Notch activation, or whether the “Polycomb” module of iRegulon merely indicates gene de-repression, has not been addressed formally in this study.

### A polarity-loss response required for NS

In order to validate the functional relevance of the transcription factors identified in the Notch-driven neoplastic growth, we then asked whether their depletion by RNAi could alter the growth and invasiveness of the NS tissue, using the *Bx>NS* fly line described previously. We first focused our analysis on the factors associated with the “oncogenic module”.

We confirmed earlier reports that blocking JNK activity by the overexpression of a JNK dominant negative construct, strongly abolishes NS driven growth (GFP positive tissue size; Fig. 4B&E) and invasiveness (Mmp1 expression; Fig. 4B&F). As shown in the *RasV12/scrib-*paradigms, RNAi mediated knock-down of the Jak/Stat pathway terminal transcription factor *stat92E*, and to a lesser extent *ftz-f1* (Atkins et al., 2016; Davie et al., 2015; Külshammer et al., 2015; Toggweiler et al., 2016) strongly suppressed both growth (GFP) and invasiveness (Mmp1; Fig. 4E&F). Unlike the RasV12 models, we did not identify in the different iRegulon analyses of the N, S, and NS transcriptomes, any particular enrichment for Sd/TEAD, the transcriptional factor mediating the effect of Yki and of the Hippo pathway mediated growth in wing discs. However, impairing Yki activity (through RNAi-mediated knock-down) strongly suppressed NS neoplastic behaviors (Fig. 4C,E&F).

**Figure 4.**
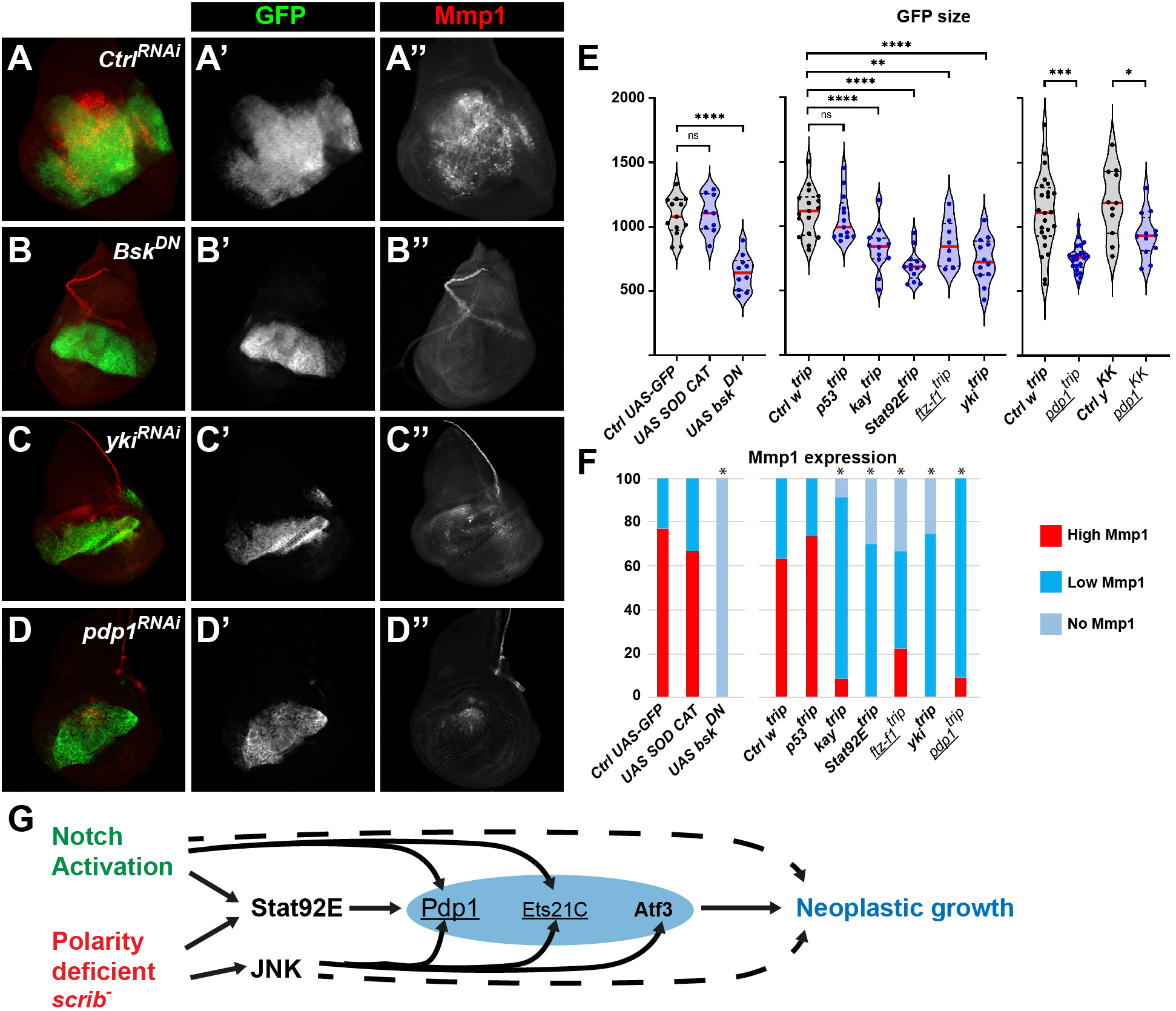
Neoplastic growth is mediated by an AP-1/Stat/Yki/Ftz-f1 module and by a diverse network of bZIP transcription factors including Pdp1. **A-D.** 3rd instar wing imaginal discs expressing *GFP*, an activated form of Notch (*Nicd*) and an RNAi for *scrib* under the control of the *Bx-Gal4* driver (dorsal wing pouch) and stained for GFP (green, white A’-D’) to assess tissue overgrowth and for Mmp1 (red, white A’’-D’’) to assess tissue invasiveness. Discs also expressed under *Bx-Gal4* control the either *UAS Bsk DN* (B) or the indicated RNAi constructs: *w[HMS00045]* (A; Ctrl), *yki[HMS00041]* (C), *Pdp1[HMS02030]* (D). Representative discs are shown. **E.** Quantification of the overgrowth of the GFP territory in the indicated genotypes. Results are shown as violin plots of total GFP area measured in arbitrary pixels. In grey are represented the controls for each experiment. Statistical tests: ANOVA (left and middle graphs) or unpaired two tailed t-test (right graph); n.s. non significant; * p<0.05; ** p<0.01; *** p<0.001; **** p<0.0001. For more details of RNAi lines either from the TRiP collection (labelled with trip superscript) or from the Vienna collection (labelled with KK superscript) are provided in supplemental information. **F.** Quantification of the Mmp1 intensity in the indicated genotypes. Results are shown as percentage where discs were classified to fall in three categories: High Mmp1 staining intensity (red, similar to that shown in A’’), Low Mmp1 staining intensity (blue, similar to that shown in D’’), No Mmp1 staining (light blue, similar to that shown in B’’). Statistical test: chi-square test for trend or Fisher’s exact test; * p<0.05. **G.** Feedforward model of the cooperation between Notch and polarity loss during wing imaginal disc neoplastic growth. **E-G.** The Pdp1, Ets21C, and Ftz-f1 NDTs are highlighted with an underline.

Taken together these results suggest that independently of the oncogenic driver, Ras or Notch, relatively similar tumorous transcriptional networks (AP-1/Yki/Stat/Ftz-f1) are put in place during their cooperation with polarity loss (Atkins et al., 2016; Davie et al., 2015; Hamaratoglu and Atkins, 2020; Külshammer et al., 2015). Given that these nodes were also identified in the S transcriptome, we suggest that they represent a polarity loss module cooperating with oncogenic signaling pathways Ras or Notch, and likely other pathways such as Hh as was initially reported (Brumby and Richardson, 2003).

### The PAR domain containing transcription factor Pdp1 is required for neoplastic growth

A striking feature of the iRegulon analyses was the NS/S JNK module, which contained classic basic leucine zipper (bZIP) transcription factors such as Jun or Fos. These factors are involved in complex homo and hetero-dimers mediating cellular responses to different stresses such as DNA damage, oxidative stress… (Reinke et al., 2013), and their binding specificities remain difficult to tease apart. Atf3 has recently been shown to control the expression of genes involved in the maintenance of epithelial polarity, and to be specifically activated in polarity deficient cells and required in the *RasV12/scrib-*overgrowth model (Atkins et al., 2016; Donohoe et al., 2018). Strikingly, amongst the different bZIP predicted by iRegulon to control the NS transcriptome, *Pdp1* is a direct Notch target (common in N, S, and NS; Supplemental Table S3 & Fig. S4A), raising the interesting prospect that Pdp1 could act as feedforward factor to promote neoplastic growth downstream of Notch.

Knocking down *Pdp1* using two independent RNAi lines, using the *Bx>NS* fly line, led to a robust reduction of tissue growth (as measured by total GFP area; Fig. 4D&E), and to a reduction in the intensity of Mmp1 staining (Fig. 4D&F). Pdp1 (the homologue of Hepatic Leukemia Factor -HLF) has previously been linked to mitotic cell cycle and growth (Reddy et al., 2006), and shown in the *RasV12/scrib-*paradigms to have only modest effects on invasion, but none on growth. In the context of Notch (*Nicd/scrib-*), the role of *Pdp1* appeared thus more essential, probably because *Pdp1* is a direct Notch target. But while *Pdp1* was required, was it sufficient to promote NS-like tumors? Overexpression of Pdp1 was sufficient to induce cell delamination and spreading, as evidenced by the increased Mmp1 staining and the spreading of GFP positive cells on the anterior part of the disc (Fig. S4C). Strikingly, the combined overexpressions of Nicd and Pdp1 resulted in high Mmp1 staining and a much wider GFP expression domain (compared to Nicd or Pdp1 alone) indicative of invasive and/or delaminating cells extending anteriorly (Fig. S4C-E). Even though these features reproduce in part some of the NS behaviors, in particular the increased Mmp1 staining, it should be noted that the discs overexpressing both Nicd and Pdp1 appeared to grow poorly, unlike NS discs. Furthermore the Mmp1 upregulation upon Nicd&Pdp1 overexpression was not restricted to the overexpressing cells, suggesting that the combination of Nicd and Pdp1 in a restricted number of cells (under the control of the *Dpp-Gal4* driver) had non-autonomous effects both on growth and on Mmp1 expression (Fig. S4E, yellow arrowhead). Taken together, these results highlight the important role of Pdp1 in Notch-driven neoplasia (necessity), but show that the sole overexpression of Pdp1 in combination with Nicd is not sufficient for optimal growth as observed in NS tumors, suggesting that the development of NS tumors rely on specific levels of Pdp1, together with the involvement of other transcription factors (e.g. Stat92E, Ftz-f1…).

## DISCUSSION

In this study, using Notch-driven paradigms of epithelial overgrowth in *Drosophila* wing discs, we describe the molecular mechanisms underlying the cooperation between Notch and polarity loss during neoplasia. We show that epithelial polarity alterations redirect the transcriptional outcome of the Notch signaling pathway, thus defining a specific set of new neoplastic Notch direct targets. We further show that this redirection occurs mainly on pre-existing Su(H) bound regions rather than new ones. Finally, we show that similarly to what was previously described for Ras signaling (Atkins et al., 2016; Davie et al., 2015), the AP-1/Stat/Yki/Ftz-f1 transcription factors are required for the cooperation between Notch signaling and polarity loss during neoplastic growth.

While cancer genomes exhibit multiple mutations in cancer cells, their functional interactions remain difficult to monitor and model. Neoplastic tissues, generated upon the combination of Notch pathway activation and polarity loss through *scrib* mutation, experience many cellular stresses: DNA damage responses, but also ER and unfolded protein response, starvation, or oxidative stresses. However, even though present, these different stresses and in particular oxidative stress and DNA damage are not individually necessary in the context of polarity loss as blocking them or the cellular response they promote (CAT/SOD overexpression, or inhibition of p53) could not significantly suppress the NS tumorous behaviors. These observations suggest that the different stress pathways activated during polarity loss are either not required for fueling growth (they are rather a consequence than a cause of neoplastic growth), or might act “redundantly” to activate a common core response required for increased growth.

While *Drosophila* and mouse models have demonstrated that overactive signaling pathways cooperate with epithelial polarity impairment to generate neoplastic growth (Brumby and Richardson, 2003; McCaffrey et al., 2012; Pagliarini and Xu, 2003; Xue et al., 2013), the vast majority of studies seeking to understand the underlying mechanisms, have focused primarily on the cooperation between activated RasV12 and *scrib* mutants, especially in *Drosophila* (Atkins et al., 2016; Cordero et al., 2010; Davie et al., 2015; Igaki et al., 2009; Katheder et al., 2017; Pagliarini and Xu, 2003; Toggweiler et al., 2016; Wu et al., 2010). Importantly, the current study, investigating the cooperation between Notch and polarity, shows that many observations made for Ras can be extended to Notch, suggesting that the paradigms used are not a Ras specificity but might represent a more general tumor growth paradigm. But, since the main, if not only, Notch pathway outcome is transcriptional, the *Nicd/scrib-*model allowed to study in greater details the modes of cooperation. The cooperation between Notch pathway activation and polarity loss led to a specific transcriptional programme, and in particular the activation of new Notch direct targets. We could show that this was not the consequence of a general redeployment to new target genes loci of Su(H), the Notch pathway-dedicated transcription factor, ruling-out one possible model for the oncogene/polarity cooperation. What could thus be the mechanisms controlling which genes were activated in the different conditions? All “Notch” activating transcriptional complexes comprise of Nicd, Mastermind, and Su(H). While no differences in overall levels of Nicd and Su(H) could be detected by western-blot (data not shown), they could be differently modified post-translationally leading to different Notch responses (core Notch response, N-only, NS-only…). Indeed, recent reports point towards different post-translational modifications for Su(H) (Frankenreiter et al., 2021). Whether they lead to different transcriptional programmes, and whether they occur *in vivo* in the N and NS models remain however to be studied. Through the use of iRegulon, we could show that the genes of the NS transcriptome, and most importantly the NS Notch direct targets, were enriched in their regulatory regions for elements corresponding to specific transcription factors, and in particular Stat92E or bZIP factors. The fact that similar transcription factor families were found in the whole up-regulated genes, and in the more limited Notch direct genes, suggest that the Notch out-put was controlled, at least in part, by factors that act more broadly on the genome. These analyses support a model where polarity loss redirects the out-put of the Notch transcriptional programme by the action of cooperating transcription factors. However, further work such as detailed comparative ChIP analyses of the different factors in the different conditions, is required to firmly establish this model.

Even though we could highlight the involvement of a similar “oncogenic module” as identified for the *RasV12/scrib-*neoplastic model (Atkins et al., 2016; Davie et al., 2015; Külshammer et al., 2015), there are specifics that are likely oncogene specific. First, unlike what was reported for *RasV12/scrib-*transcriptomes, Yki/Sd/TEAD modules were not found enriched in the different *Notch* and *scrib-*transcriptomes. In the case of Ras, it was shown that Yki activity could reprogram Ras by promoting the expression of the Ras pathway specific regulators Capicua and Pointed to promote aggressive growth (Pascual et al., 2017). Both genes were either unaffected (*capicua*) or downregulated (*pointed*) in NS Notch driven neoplastic paradigm, suggesting that, even though Yki is clearly required (Fig. 4C), changes in the expression of capicua and pointed are unlikely mediators here. These differing results in the enrichment of Yki/Sd/TEAD motifs between Notch and Ras transcriptomes in the context of polarity loss, might reflect the inhibitory effect Notch has on Yki activity in the wing pouch, in part through the action of the *vestigial* (Djiane et al., 2014). Furthermore, in NS transcriptome, we identified a contribution of the E(spl) bHLH transcriptional repressors, canonical Notch targets (Bray, 2016), which represents thus a Notch specificity. However, the fact that motifs for E(spl)-HLH repressors are found in the up-regulated transcriptome of NS and not N could suggest that in NS, the repressive ability of E(spl)-HLH factors is antagonized, further allowing higher expression of Notch targets. More precisely, our previous work identified many incoherent feed-forward loops in the N hyperplastic transcriptome, including through the action of E(spl) repressors (Djiane et al., 2013), which might thus be resolved in NS. It would be interesting to explore further the link between NS and E(spl)-HLH-mediated repression, but due to the high redundancy between the seven E(spl)-HLH factors (δ, γ, β, 3, 5, 7, 8) and Dpn, the requirement of E(spl)-HLH-mediated repression in the Notch-driven neoplasia could not be formally tested.

While performing functional assays to identify the genes and processes required for NS tumor growth, we could show that the Notch direct targets associated with “de-novo” NS-specific Su(H) peaks were unlikely to be major contributors. We show however that the bZIP PAR domain containing factor Pdp1 is required for NS tumor growth and invasiveness. Su(H) is bound in the vicinity of Pdp1 in all wing discs set-ups and in particular in N and NS, and Pdp1 represents a “core” Notch target activated in all overgrowth conditions, albeit at higher levels in polarity deficient conditions (Fig. S4A). Pdp1 is not only a Notch target, but also a Jak/Stat target, at least in the developing eye, and canonical tandem Stat92E putative binding sites are found in its second intron, although not overlapping with Su(H) binding which is found in its first intron (Flaherty et al., 2009). Interestingly, Pdp1 is required for Stat92E phosphorylation and efficient Jak/Stat signaling (Baeg et al., 2005), suggesting that Notch might amplify Stat92E signaling during wing disc neoplastic growth, both through ligand expression (Upd ligands are Notch direct targets; this study; (Djiane et al., 2013)), and Pdp1 expression.

Even though Pdp1 downregulation could suppress NS neoplastic growth, it was not as efficient as JNK inhibition, or Yki downregulation, suggesting that other factors in parallel to Pdp1 might be involved, such as the previously identified Atf3 (Donohoe et al., 2018), but also the other Notch direct target Ets21C (Fig. 2D; (Külshammer et al., 2015; Toggweiler et al., 2016). Indeed, RNAi-mediated knock down of *Atf3* or *ets21C* partly suppressed *Bx>NS* tumor growth (GFP) and invasiveness (Mmp1; data not shown). This action of both Pdp1 and Ets21C suggest a feedforward loop downstream of Notch that in the context of polarity loss and JNK activity promotes neoplastic growth (Fig. 4G). However, since *Atf3, Pdp1*, and *Ets21C* (but also *Ftz-f1*) are all upregulated in N hyperplastic conditions, their sole upregulation cannot be sufficient for neoplasia. The fact that Atf3 and Pdp1 iRegulon enrichments are not found in N (Fig. 3 and S3), could indicate that despite being upregulated in hyperplastic N, their transcriptional activities are hindered, or that one key cooperating factor enabling their action is missing. Further studies are thus required to test this possibility and study how, in the context of normal epithelial polarity, Notch activation prevents the action of Pdp1/Ets21C/Atf3, thus preventing the transition to neoplasia.

## MATERIALS AND METHODS

### Drosophila genetics

The different overgrowth paradigms were obtained by generating random clones in 3rd instar wing discs at high frequency as previously published in (Djiane et al., 2013). In brief, the *abxUbxFLPase; Act>y>Gal4, UAS GFP; FRT82B tubGal80* flies were crossed either to *FRT82B* (to generate Ctrl discs), or to *UAS-Nicd; FRT82B* (to generate hyperplastic N discs), or to *UAS-Nicd; FRT82B scrib1* (to generate neoplastic NS discs). *scrib1* represents a loss of function allele for the *scribble* gene. Because *scrib1* clones are eliminated in growing discs, the dysplasic S discs were obtained from *FRT82B scrib1 / Df(3R)BSC752* 3rd instar larvae. All crosses were performed at 25°C and carefully staged (time after egg laying and tube crowding).

For transcriptomic and ChIP analyses, discs were dissected from these carefully timed animals. N, S, and NS overall larval body sizes were very similar to WT controls at day 5 and 6, suggesting that they grew normally during 2nd and 3rd instar larval stages. All animals with wing disc growth defects displayed pupariation delay: S and NS never pupate, while some (not all) N animals eventually pupate at day 11 or 12 after egg laying (ael; normal time being 5 to 6 ael). These observations suggest that the main timing problem for N, S, and NS was in the transition to pupa at 5 to 6 ael. By monitoring differences at day 6 ael, just after spiracle eversion (a classic landmark in developmental timing), we thus monitored differences just prior to the extension of larval life, and very far from the extreme that could be observed, suggesting that tissues would remain comparable.

For functional studies, neoplastic growth was obtained by driving *UAS-Nicd* and the *scrib* RNAi *P{TRiP.HMS01490}attP2* by the *Bx-Gal4* (pouch of larval wing discs). Modifications of the overgrowth phenotype and of the expression of the Mmp1 invasive marker were performed by crossing in F1 *Bx-Gal4, UAS GFP;; UAS Nicd, UAS scribHMS01490* to the desired *UAS RNAi* or control lines (*UAS white RNAi, UAS yellow RNAi*, or *UAS GFP*), to ensure similar UAS load. List of lines tested in Supplemental Materials.

Information on gene models and functions, and on *Drosophila* lines available were obtained from FlyBase (flybase.org – (Thurmond et al., 2019)).

### RNA extraction and RNA-Seq

RNA from 60 or 80 dissected third instar larva wing discs of WT, N, NS and S discs was extracted using TriZOL. Genomic DNA was eliminated using Ambion’s DNA-free kit (#AM1906). cDNA bank preparation were then performed from 1μg of RNA and sequencing on a Illumina HisSeq 2000 by the Biocampus genomic facility MGX of Montpellier. After sequencing, reads obtained were filtered based on their quality (circa 40 millions reads were kept per conditions). The reads were then align on *Drosophila* dm6 genome by the ABIC facility in Montpellier producing a matrix of reads per gene and per condition. This matrix was then normalized and pair-wise differential expression was performed using DESeq (Anders and Huber, 2010). Other differential expression tools were tested such as DESeq2 and edgeR with default parameters but appeared either less stringent, or inadequate.

### qPCR

qPCR was performed on biological triplicates on a Roche LightCycler 480, and fold change was estimated by the δδCT approach. List of primers used in Supplemental Materials.

### Su(H) Chromatin Immuno Precipitation

After dissection in PBS 1X, Protein/DNA complexes from 60 wing discs (80 for S condition) were cross-linked with 1% formaldehyde for 10 minutes. The reaction was then quenched by 0.125 M Glycine and washed 3x in PBS. Wing disc cells were resuspended in 50μL Nuclear Lysis Buffer (Tris-HCl pH 8.1 20mM, EDTA 10mM, SDS 1%). Lysates were sonicated on a Bioruptor (Diagenode), and diluted 10x in Immunoprecipitation Dilution Buffer (Tris-HCl pH 8.1 20mM, EDTA 2mM, SDS 0.01%, NaCl 150mM, Triton X-100 1%) and precleared with rabbit IgG (Sigma) and protein G Agarose (Santa Cruz Biotechnology). ChIP reactions were performed by incubating lysates overnight at 4°C with 1ng of Goat anti-Su(H) (Santa Cruz Biotechnology, sc15813), and immunocomplexes were then isolated with Protein G Agarose for 2h, washed 2x with Wash Buffer 1 (Tris-HCl pH 8.1 20mM, EDTA 2mM, SDS 0.1%, NaCl 50mM, Triton X-100 1%) and 2x with Wash Buffer 2 (Tris-HCl pH 8.1 10mM, EDTA 1mM, LiCl 250mM, NP-40 1%, Deoxycholic acid 0.4%), before a decross-linking step at 65°C in 0.25M NaCl. Samples were then treated with 0.2 mg/mL proteinase K and 50mg/mL RNase A. The DNA was then purified on columns (Qiagen, 28106). ChIP efficiency was checked by qPCR normalized on input chromatin with the following primer couples, corresponding to known strong binding sites of Su(H). List of primers used in Supplemental Materials.

For whole-genome analysis, 1 μg double-stranded ChIP or input DNA (corresponding to 180 discs for each replicate) was labelled with either Cy3-or Cy5-random primers using the Nimblegen Dual Colour kit. Both ChIP and input were co-hybridised to NimbleGen D. melanogaster ChIP-chip 2.1 M whole-genome tiling arrays in the NimbleGen hybridisation station at 42°C for 16 h and then washed according to the NimbleGen Wash Buffer kit instructions. The data obtained were normalized using quantile normalization across the replicate arrays in R. Window smoothing and peak calling were performed using the Bioconductor package Ringo (Toedling et al., 2007) with a winHalfSize of 300 bp and min.probes = 5. Probe levels were then assigned P-values based on the normalNull method, corrected for multiple testing using the Hochberg–Benjamini algorithm and then condensed into regions using distCutOff of 200 bp.

In order to determine the Notch Direct Targets (NDTs), ChIP and RNA-Seq results were compared: NDTs are defined as up-regulated genes with Su(H) enrichment within 20kb. As such one Su(H) peak could be assigned to several upregulated genes consistent with its role in enhancer regions. The 20kb window was chosen as it allowed the recovery of more than 85% of NDTs in our previous study that was based on closest gene assignment irrespective of distance (Djiane et al., 2013).

### GO Term analyses

The lists of significantly regulated genes in the various comparisons were submitted to gene ontology (GO) term enrichment analysis. We used the GO biological process (GOBP) ontology and applied hypergeometric tests (p-values) followed by Benjamini-Hochberg multiple hypothesis correction (q-values).

### iRegulon analyses

In order to determine the likely transcriptional modules in our transcriptomic and NDTs datasets, we used the online tool iRegulon (http://iregulon.aertslab.org/) (Janky et al., 2014; Verfaillie et al., 2015), with the standard settings using the 6K Motif collection (6383 PWMs) and a Putative regulatory region of “10kb upstream, full transcript and 10kb downstream”. Importantly, these settings allowed the recovery of the “positive control” Su(H) module.

### Immunofluorescence

Antibody staining of wing imaginal discs were performed using standard protocols. Briefly, larval heads containing the imaginal discs (LH) were dissected in cold PBS and fixed for 20min in 4% Formaldehyde in PBS at room temperature (RT), before being rinsed 3x 10min in PBS 0.2% TritonX100 (PBT), and blocked in PBT + 0.5% BSA (PBTB) for 30min at RT. LH were then incubated overnight at 4°C with primary antibodies in PBTB. LH were then rinsed 3x 10min in PBT at RT and before being incubated with secondary antibody in PBTB for 90min at RT. LH were then rinsed 3x 20min in PBT at RT, before being equilibrated overnight in Citifluor mounting media (Agar). Discs were then further dissected and mounted. Images were acquired on a Zeiss Apotome2 or Leica Thunder microscope and processed and quantified using Zen, Las X, or ImageJ. Primary antibodies used were rabbit anti cleaved Drosophila Dcp-1 (9578, Cell Signaling Technologies, 1:200), rat anti-DE-Cadherin (DCAD2, Developmental Studies Hybridoma Bank – DHSB, 1:25), rabbit anti-GFP (A6455, Molecular Probes, 1:200), and mouse anti-Mmp1 (3A6B4, DHSB, 1:25). Secondary antibodies used conjugated to Alexa-350, Alexa-488, or Cy3 were from Jackson Labs Immuno Research (1:200).

### Quantification methods

Genotypes were tested in batches with controls and 10-20 images corresponding to 10-20 different discs were all acquired on the same microscope with the same exposure settings.

Growth was estimated by the size of the GFP positive area measured as pixel numbers. Mmp1 intensities were ranked as High, Low, Null by independent observer with genotypes masked and processed in random order. Graphs and statistics (indicated in the legends) were performed using the GraphPad Prism software.

### Raw data

The data discussed in this publication have been deposited in NCBI’s Gene Expression Omnibus (Edgar et al., 2002) and are accessible through GEO Series accession number GSE185339 (https://www.ncbi.nlm.nih.gov/geo/query/acc.cgi?acc=GSE185339), and GSE185490 (https://www.ncbi.nlm.nih.gov/geo/query/acc.cgi?acc=GSE185490).”

## ACKNOWLEDGEMENTS

We thank G. Alvès, D. Andrew, D. Eberl, P. Léopold, R. Levayer, C. Maurange, A. Teleman, J. Terman, and T.T. Su for sharing flies. We acknowledge the Bloomington Stock Center, the Vienna Stock Center, the DGRC Kyoto Stock Center, the Developmental Studies Hybridoma Bank, the *Drosophila* facility, the MGX sequencing facility (BioCampus Montpellier, CNRS, INSERM, Université de Montpellier), and FlyBase for their support to our research.

The lab of SJB is supported by the MRC (MRC programme grant MR/L007177/1). RL was supported by fellowships from “Ligue Nationale Contre le Cancer - LNCC” and from “Fondation ARC pour la recherche sur le cancer”. The lab of AD is supported by the “Fondation ARC pour la recherche sur le cancer”, “Marie Curie CIG”, and “Agence Nationale de la Recherche - ANR-18-CE14-0041”.

## AUTHOR CONTRIBUTIONS

RL, CG, PL, MRV, DK, and AD performed experiments; RL, CG, LHM, JC, and AD analyzed data; JC provided expertise for RNA-Seq; BF and SJB provided technical support and expertise for ChIP; RL, SJB, and AD wrote the manuscript; AD conceived and supervised the project.

## SUPPLEMENTAL INFORMATION

### SUPPLEMENTAL MATERIAL AND METHODS

#### Drosophila genetics

Driver lines were *Bx-Gal4* (wing disc pouch), *Dpp-Gal4*, and *Ptc-Gal4* (both wing disc antero-posterior boundary). Overexpression lines were *UAS Nicd* (made by the Bray lab), *UAS GFP:Act87E [7-6]* BL#9249, and *UAS Pdp1.T* BL#78087.

Overexpression lines tested in the *Bx-Gal4, UAS GFP;; UAS Nicd, UAS scribHMS01490* screen were *UAS GFP*, *UAS bskK53R [20.1a]* BL#9311, and *UAS SOD CAT* (gift from P. Leopold). RNAi lines used are listed in the table below with an indication of the labels used in Fig. 4E&F. TRiP collection lines have a stock BL#, and Vienna collection lines have a stock v#. While performing the *Bx>NS* modifier experiments, we used controls originating from the same collection: *RNAi white* for TRiP lines, and *RNAi yellow* for Vienna KK lines.

| Gene | RNAi ID | Stock # |
| --- | --- | --- |
| Act87E | HMS02488 | BL#42642 |
| Act87E | KK111781 | v#102480 |
| cdi | KK100725 | v#109409 |
| CG7059 | GD10763 | v#21651 |
| CG13850 | KK104804 | v#100863 |
| CG13857 | GD6226 | v#44061 |
| CG18596 | KK104660 | v#108183 |
| ftz-f1 | HMS00019 | BL#33625 |
| Ir87a | HMJ22848 | BL#60476 |
| Ir87a | KK106593 | v#100667 |
| kay | HMS00254 | BL#33379 |
| p53 | HMS02286 | BL#41720 |
| Pdp1 | HMS02030 | BL#40863 |
| Pdp1 | KK109014 | v#110551 |
| REG | KK102083 | v#110156 |
| stat92E | HMS00035 | BL#33637 |
| vito | KK111866 | v#102513 |
| w | HMS00045 | BL#33644 |
| y | KK104196 | v#106068 |
| yellow-e | HMC06250 | BL#65970 |
| yellow-e | KK106243 | v#100926 |
| yellow-e3 | KK106158 | v#105879 |
| yki | HMS00041 | BL#34067 |

#### Primers

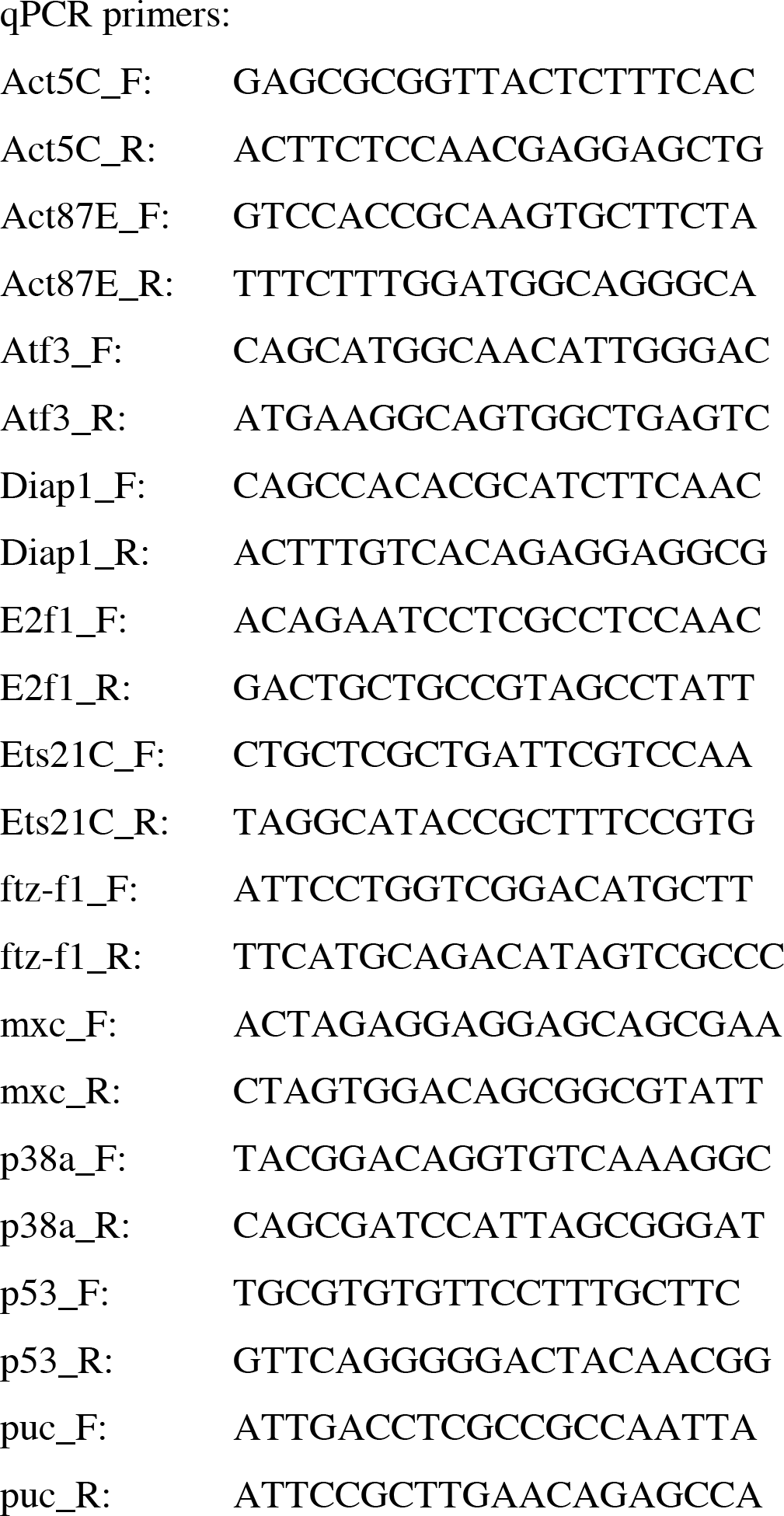
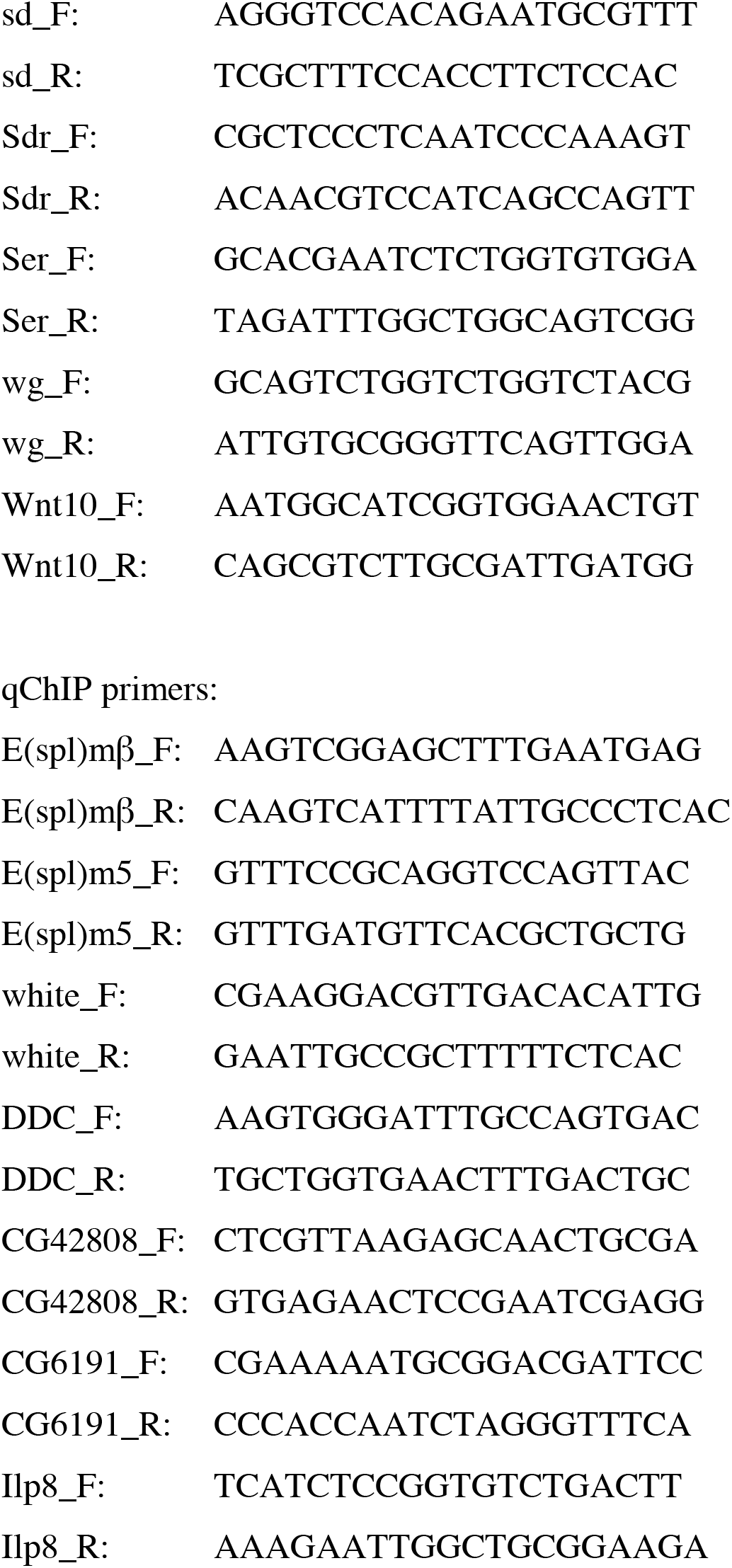

### SUPPLEMENTAL FIGURE LEGENDS

**Figure S1.**
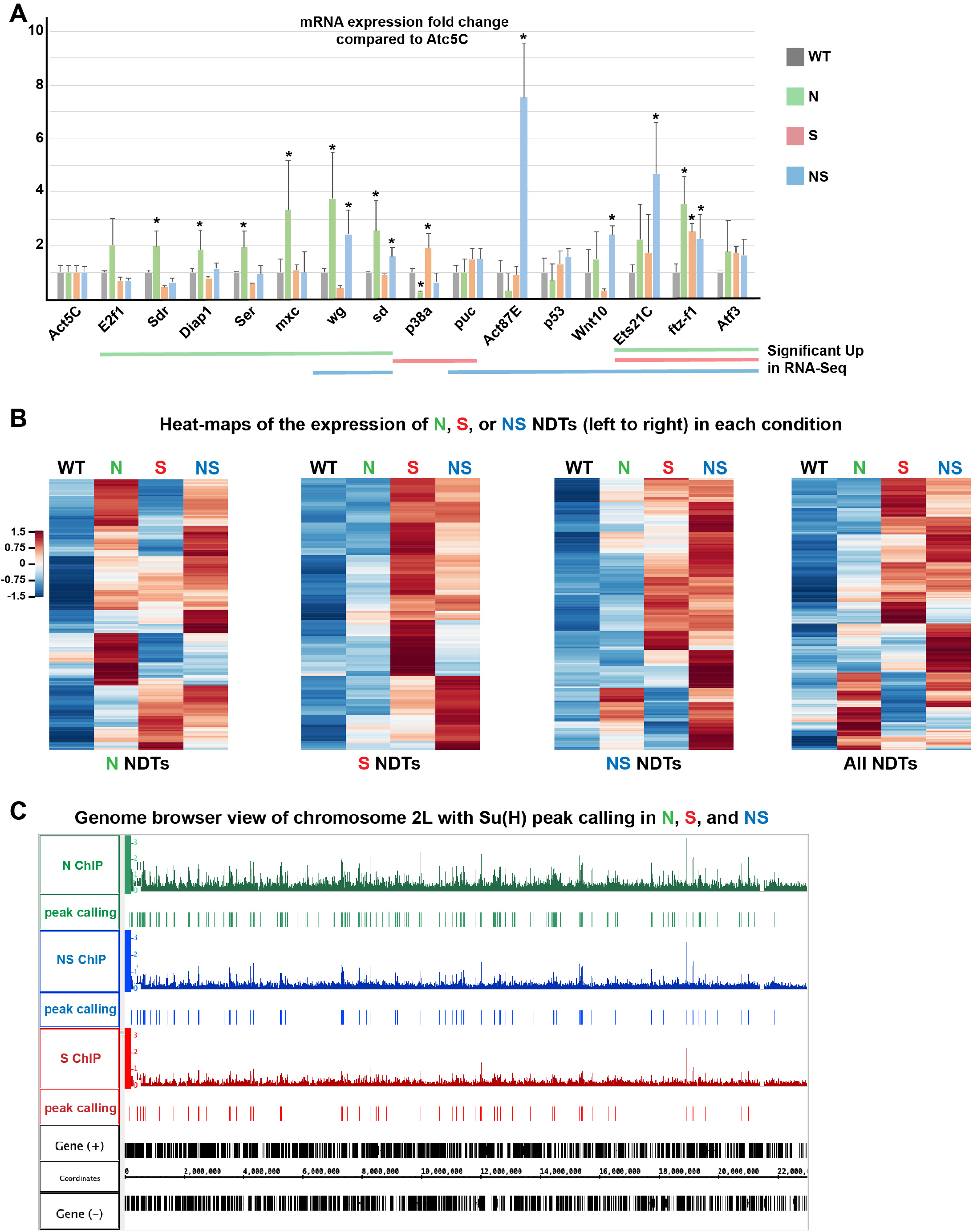
Features of the Notch Direct Targets (NDTs) in N, S, and NS (relates to Fig. 3) **A.** Semi-quantitative RT-PCR of the indicated genes represented as fold change compared to WT (grey) in the different N (green), S (red), and NS (blue) growth paradigms and normalized to *Atc5C* expression. Biological triplicates, standard error to the mean (s.e.m.) is shown. ANOVA statistical test, * p<0.05. **B.** Heatmaps for the expression of the different NDTs in WT, N, S, and NS. From left to right are presented the N, S, NS, and finally All NDTs, highlighting that NDTs could be transcriptionally up-regulated in more than in one condition. **C.** Genome browser view of the whole left arm of the 2nd chromosome, and showing the Su(H) ChIP enrichment (upper rows) and the intervals called as Su(H) peaks (lower rows) in N (green), NS (blue), and S (red). Note the higher number of peaks in N, and the rarity of NS, or S peaks not found in N.

**Figure S2.**
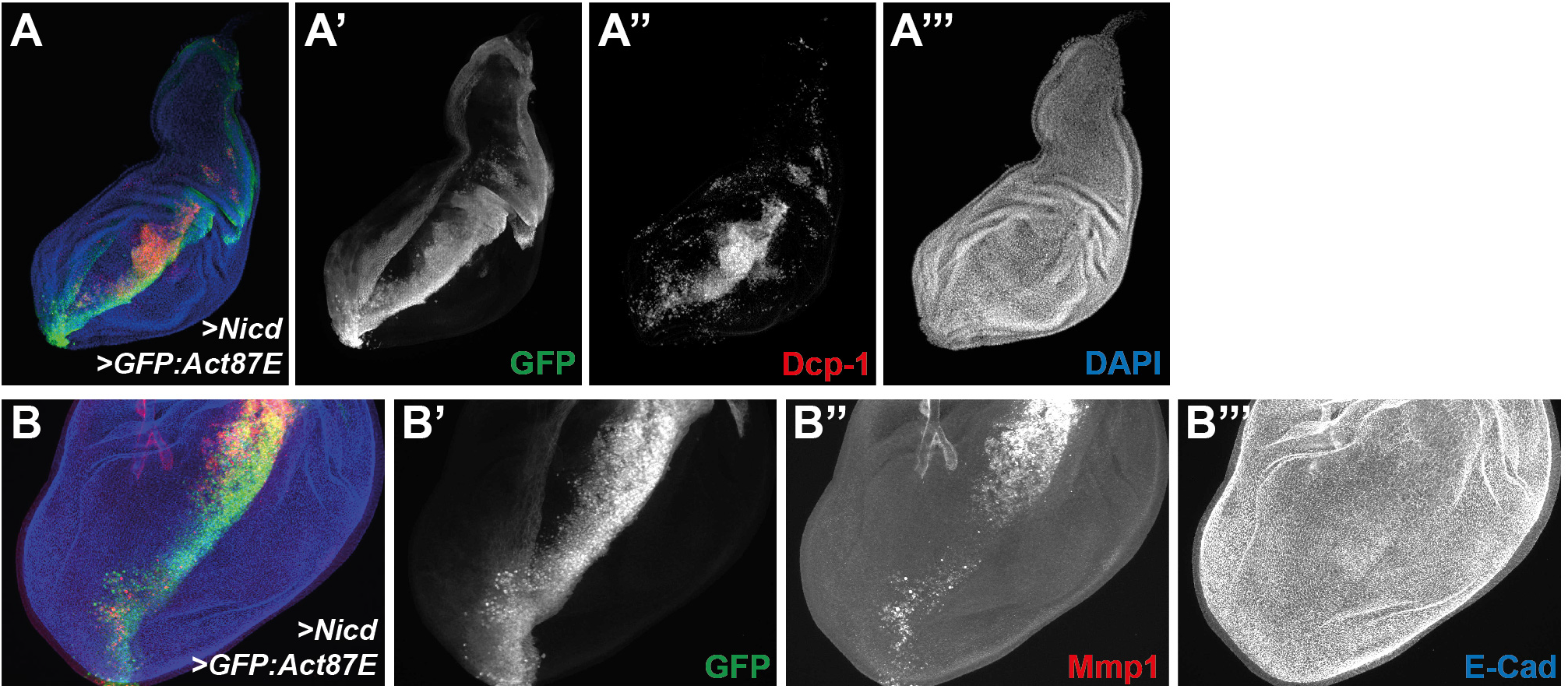
Act87E promotes Mmp1 expression and cell delamination. **A-B.** GFP:Act87E overexpressed together with activated Notch (Nicd) under the control of the *Ptc-Gal4* driver (antero/posterior boundary cells in green, A’&B’), promotes the expression of the Dcp-1 caspase (red, A’’), and the metalloprotease Mmp1 (red, B’’). Similar results were obtained for the sole overexpression of GFP:Act87E (without Nicd). DAPI (blue, A’’’) or E-Cad (blue, B’’’) mark all wing disc cells. (A) Whole wing disc. (B) Detail of the overgrowing wing pouch (magnification in B is twice that in A).

**Figure S3.**
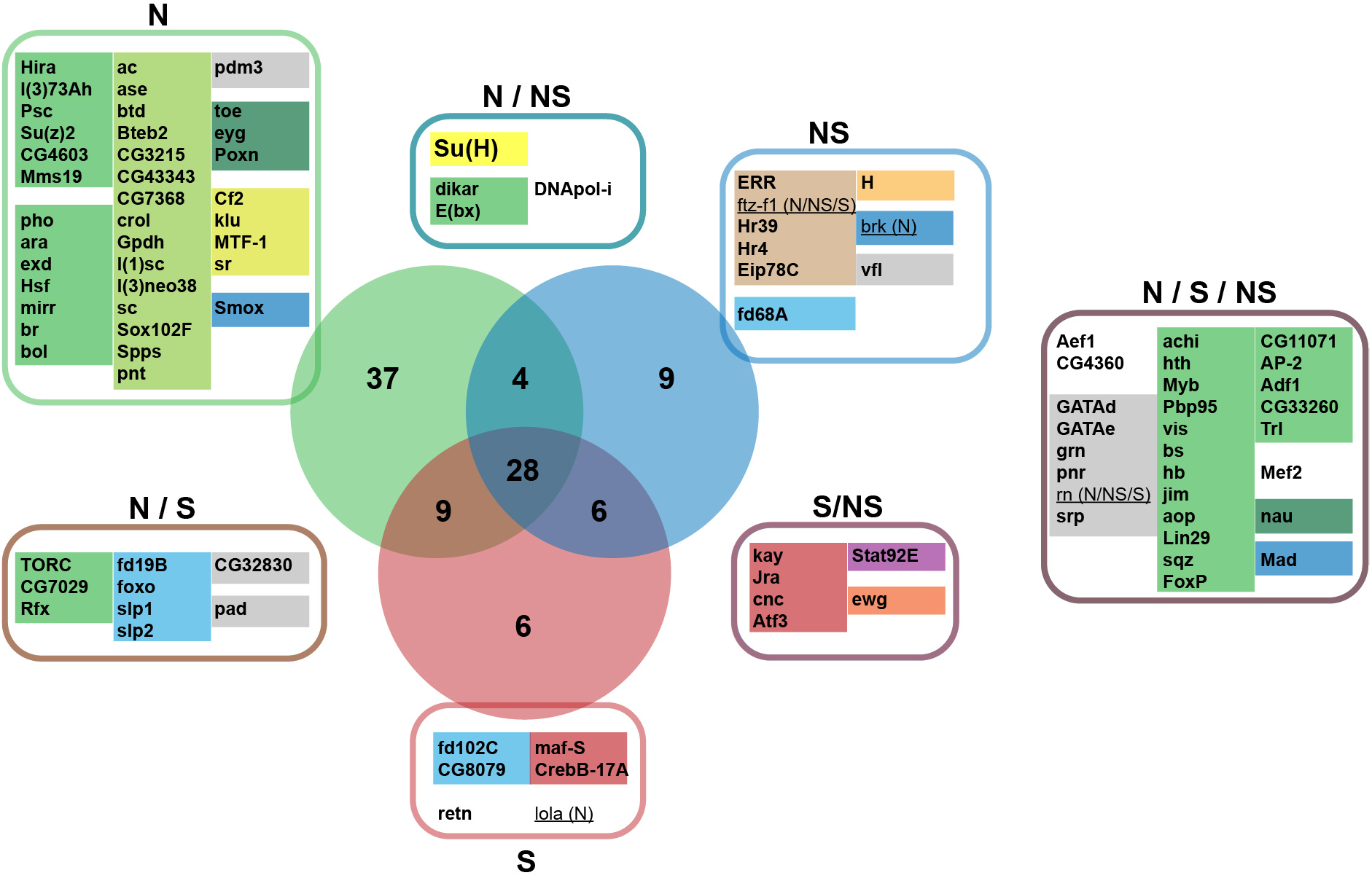
Identification of potential transcriptional modules mediating N, S, and NS growth (relates to Fig. 3) Venn diagram of significant transcription factors (TFs) identified by iRegulon as potential key mediators for the expression of the N (in the green circle), S (red circle), and NS (blue circle) Notch Direct Target (NDT) genes. Fed with lists of co-regulated genes, and analyzing the genomic features in the vicinity of the transcription start sites of these genes, iRegulon identifies potential groups of TFs and DNA-binding factors, that are enriched in the dataset of regulatory sequences, and could thus represent potential mediators regulating the expression of NDT genes in N, S, and NS. For each condition, TFs were identified as part of “regulons” or group of TFs that could potentially together regulate the expression of subsets of the transcriptome. Taking only the TFs corresponding to the significant regulons identified in N, S, and NS, the shared and unique potentially regulating TFs are here presented in the different colored boxes. In each box, TFs were then grouped according to their molecular family (suggesting pretty similar binding motifs on the DNA), and color-coded: for instance, bZIP TFs in light maroon, or the nuclear receptors in brown. The molecular family color code was not respected for the green groups which correspond to very diverse “regulons” that appear linked to epigenetic chromatin regulators/remodelers. For the complete list of regulons identified in each condition, and the nature of TFs and DNA binding proteins in each regulon, see Supplemental table S5). Numbers in the Venn diagram represent the number of TFs identified. TFs that are also NDTs and which could participate in feedforward loops are underlined (with their NDT condition in parentheses).

**Figure S4.**
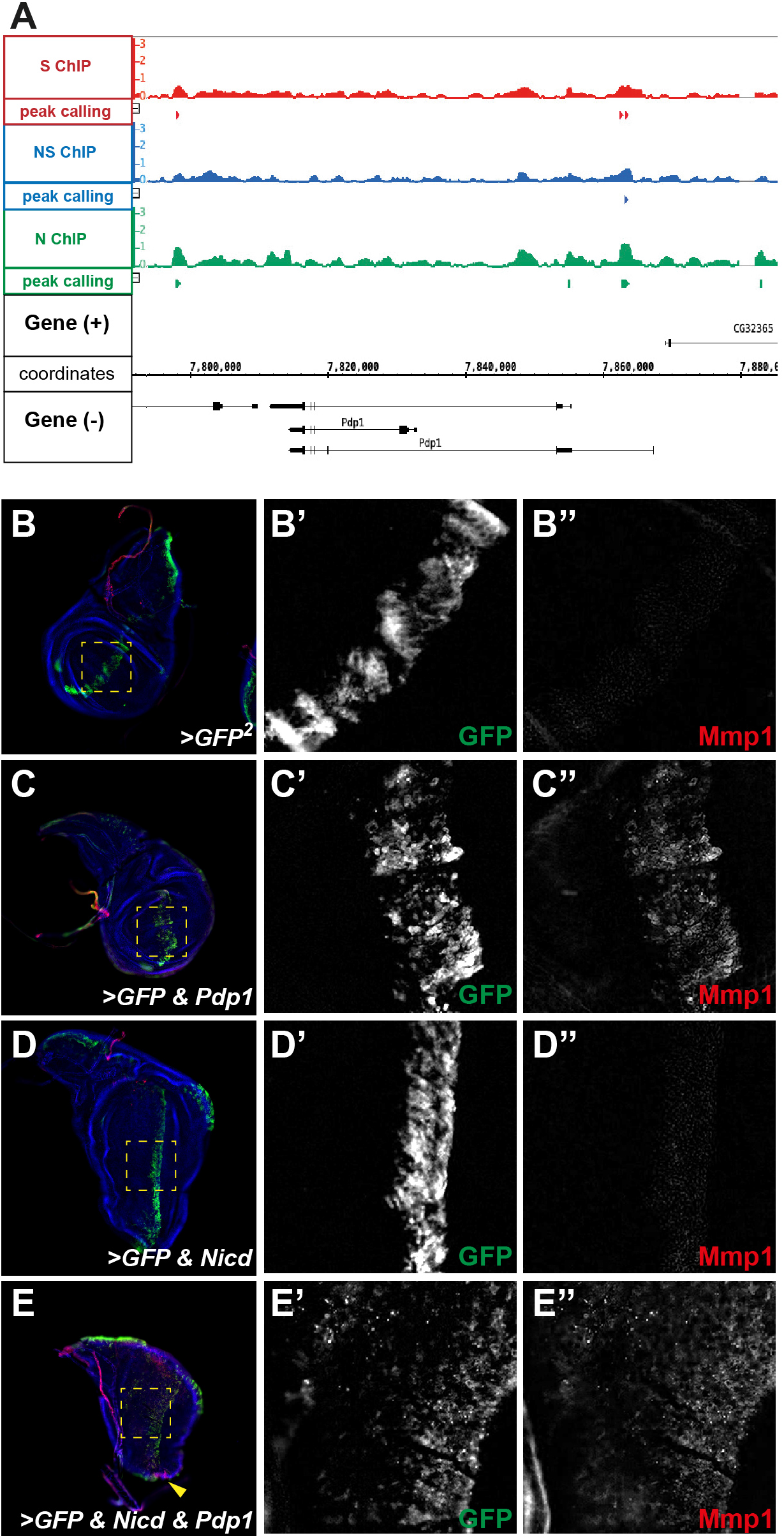
Pdp1 is a direct Notch target (relates to Fig. 4) **A.** Genome browser view of the *Pdp1* locus, and showing the Su(H) ChIP enrichment (upper rows) and the intervals called as Su(H) peaks (lower rows) in N (green), NS (blue), and S (red). **B-E.** Pdp1 overexpression causes Mmp1 expression. GFP alone (B) or in combination with *Pdp1* (C), with *Nicd* (D), or with *Nicd & Pdp1* (E) was overexpressed using the *Dpp-Gal4* driver. Higher magnification corresponding to the yellow dashed boxes show GFP in green (B’-E’) and Mmp1 in red (B’’-E’’). E-Cad used as landmark is shown in blue (B-E). Pdp1 overexpression resulted in Mmp1 positive cells extending anteriorly (C). This was enhanced when combined with Nicd (E). Representative discs shown (out of 18 imaged discs from 3 experiments). **(E)** Mmp1 expression is found in the *Pdp1&Nicd* overexpressing cells (GFP positive under the influence of the *Dpp-Gal4* driver), but also in non-expressing cells (non-autonomous, yellow arrowhead).

### SUPPLEMENTAL TABLES

**Table S1.** Differentially expressed genes in N, S, and NS identified by DESeq (related to Fig. 1). Columns are:

FBgn_ID: Unique FlyBase gene ID

Symbol: Current FlyBase gene symbol

qval: adjusted p-value for multiple testing

logFC: log2 of the Fold Change “Condition N, S, or NS” / “Control WT”

**Table S2.** Su(H) ChIP enrichment peaks coordinates in N, S, and NS (related to Fig. 3). Columns are:

Exp: N, S, or NS

Chr: Chromosome arm

MIN: smallest peak coordinate

MAX: biggest peak coordinate

**Table S3.** All Notch Direct Targets (NDTs) ordered by genomic position. This table includes an indication whether the genes are transcriptionally upregulated or have a Su(H) peak in the vicinity in each N, S, and NS condition. Columns are:

N/NS/S: NDT in the corresponding condition

Type: NDT in different conditions.

FBgn_ID: Unique FlyBase gene ID

SYMBOL: Current FlyBase gene symbol

K_ARM: Chromosome arm location of the gene

MIN (gene pos): smallest gene coordinate

MAX (gene pos): biggest gene coordinate

STRAND: +1 or −1

N Fold: Log2 Fold Change in gene expression N/WT (n.s. not significant)

N ChIP: Su(H) ChIP enrichment peak within 20kb in N (green yes, red no)

NS Fold: Log2 Fold Change in gene expression NS/WT (n.s. not significant)

NS ChIP: Su(H) ChIP enrichment peak within 20kb in NS (green yes, red no)

S Fold: Log2 Fold Change in gene expression S/WT (n.s. not significant)

S ChIP: Su(H) ChIP enrichment peak within 20kb in S (green yes, red no)

**Table S4.** Curated iRegulon analyses of the significantly upregulated genes in N, S, and NS (related to Fig. 3). Analyses were performed using the 6K-PWM and 10kb upstream and downstream set-ups.

**Table S5.** Curated iRegulon analyses of the Notch Direct Targets in N, S, and NS (related to Fig. S2). Analyses were performed using the 6K-PWM and 10kb upstream and downstream set-ups.

